# Open-access template and database approaches for pseudo-CT generation in brain PET/MRI attenuation correction

**DOI:** 10.64898/2025.12.22.695942

**Authors:** Christian Milz, Matej Murgaš, Inés Merida, Leo R Silberbauer, Lukas Nics, Godber M Godbersen, Gregor Gryglewski, Marcus Hacker, Nicolas Costes, Alexander Hammers, Rupert Lanzenberger, Andreas Hahn, Murray B Reed

## Abstract

Positron emission tomography (PET) provides high-sensitivity molecular information in neuroimaging. However, its quantitative accuracy critically depends on attenuation correction (AC). Unlike PET/CT, hybrid PET/MRI systems cannot directly measure photon attenuation, necessitating alternative AC strategies. While several MRI-based AC tools exist, they often raise concerns regarding data privacy, long-term accessibility and reproducibility. We present an open-access framework for PET AC using pseudo-CT (pCT) methods. A database of 60 paired MRI-CT scans was curated and normalized to stereotactic MNI space. Three established pCT approaches (Boston, MaxProb, UCL) were adapted to enable their use with the normalized database. Method performance was assessed by comparison with a CT-based AC reference in an independent validation sample of 28 subjects, using the Jaccard index and [^11^C]DASB PET-derived serotonin transporter quantification as evaluation metrics. Boston and MaxProb demonstrated consistent performance across database subsets, while the UCL achieved the closest agreement with CT-derived AC and yielded the lowest quantification error (mean relative error <0.5%). This work demonstrates the feasibility of different pCT approaches using a database in MNI space thereby introducing an openly available reference database to support fast, reproducible and continued integration of MRI-based AC into PET reconstruction pipelines.

## INTRODUCTION

Positron emission tomography (PET) is an essential tool for studying both normal brain function and pathological alterations across diverse patient populations. The field of molecular imaging with PET now spans a broad range of radioligands, outcome measures from standardized uptake value (SUV) to volumes of distributions (1) and more recently, stimulation-specific activation (2–4).

Accurate reconstruction of radioligand concentration is critical for all these approaches, and a key feature thereof is attenuation correction (AC), which requires an estimate of photon attenuation in the head (5). Traditionally, AC is performed using transmission scans with e.g., a ^68^Ge source rotated around the subject, or more recently X-ray computed tomography (CT) (6,7). However, in hybrid PET/MRI systems these reference standards are not available without additional scans, and standard MRI sequences cannot fully characterize tissue density.

To address this limitation, alternative methods have been developed to create so-called pseudo-CTs (pCTs) for AC in PET/MR imaging. These fall into three broad categories: segmentation-based, database-based and emission-based AC. Segmentation-based AC (8–10) identifies tissue types utilizing specialized MRI sequences. Database AC approaches (11–14) align pairs of MRI and CT from a database to the subject’s anatomy. Emission-based AC estimates attenuation directly from PET data using, for example, maximum likelihood algorithms (15) or deep learning methods (16). An in-depth comparison of various attenuation correction methods exceeds the scope of this work, and readers are referred to other publications (5,6,17–21).

Beyond accuracy, practical factors such as accessibility, ease of integration and adaptability to evolving research needs are increasingly important aspects of software tools. The neuroimaging community has embraced these principles throughout widely used open frameworks such as SPM12 (https://www.fil.ion.ucl.ac.uk/spm/), AFNI (https://afni.nimh.nih.gov/) and FSL (https://fsl.fmrib.ox.ac.uk/fsl/fslwiki/) for processing of imaging data, data management with Brain Imaging Data Structure (BIDS) (22), anatomical atlases (23) as well as new approaches to kinetic modelling (24) and functional PET (25).

With respect to AC, several database pCT generation tools (e.g.: UCL (http://niftyweb.cs.ucl.ac.uk/); (11)); MaxProb (13,26)) are available for online use. Notably the RESOLUTE algorithm (10) is available on GitHub for local installation, but was found to be sensitive to acquisition parameters and scanner software version (14). Online tools enable creation of pCTs with minimal computational resources on the user’s end but raise potential data protection concerns, as they require uploading unprocessed (i.e., non-defaced, native space), individual T1-weighted images. In addition, processing time can be long since the entire database must be registered to each individual T1-weighted image. Requests from several users at the same time might lead to longer processing queues, increasing waiting time. Moreover, maintenance or long-term availability are not guaranteed and custom solutions are not supported.

In this work, we present adaptations of the MaxProb, Boston and UCL database approaches for pCT creation, using a database that is pre-aligned to MNI space. This strategy aims to improve computational efficiency, avoids sharing raw subject data, and facilitates seamless integration into existing PET/MRI reconstruction pipelines. We further examine how database size influences pCT performance, which is evaluated with an independent validation sample of subjects acquired within the same study but not included in the database. To support accessibility, we provide an openly available database of MRI-CT pairs, corresponding templates and a lightweight MATLAB implementation for use with the Siemens reconstruction software.

## MATERIALS and METHODS

### Subjects and study design

In total, 88 subjects (mean age ± SD = 27.5 ± 8.7, 52 female) previously included in a clinical trial (EudraCT Number: 2014-001550-42) were selected for creating pCTs (n = 60) and validation of pCTs for attenuation correction (n = 28). More detailed information on the subjects used for the various templates and validation can be found in Table 1. Part of the sample was already published previously in a different context (27–29).

**Table 1:** Demographic information on database subsets and validation set. Subjects in the validation set are not included in the database.

| | n | Sex (f/m) | mean Age ± SD | HC/MDD |
| --- | --- | --- | --- | --- |
| db5 | 5 | 3/2 | 25.2 ± 7.3 | 3/2 |
| db10 | 10 | 5/5 | 27.6 ± 5.5 | 8/2 |
| db20 | 20 | 12/8 | 25.8 ± 7.5 | 14/6 |
| db40 | 40 | 23/17 | 26.0 ± 6.5 | 25/15 |
| db60 | 60 | 36/24 | 25.4 ± 7.5 | 38/22 |
| validation set | 28 | 16/12 | 32.1 ± 11.6 | 13/15 |

Subjects underwent general medical examination at the screening visit, including routine laboratory tests, electrocardiography, and neurological assessment. Exclusion criteria included a history of neurological disorders, substance abuse, smoking, pregnancy, or current breastfeeding. Alongside healthy volunteers (n = 51), subjects diagnosed with major depressive disorder (MDD) (n = 37) were included in the analysis. The study (clinical trial including attenuation correction methodology) was approved by the Ethics Committee of the Medical University of Vienna (1307/2014) and procedures were carried out in accordance with the Declaration 1964 of Helsinki. All subjects gave written informed consent. Sharing of the MR-CT image pairs was approved by departments of legal affairs and data clearing of the Medical University of Vienna (2025–042).

### Neuroimaging

All subjects underwent a low-dose CT scan using a Biograph TruePoint PET/CT (Siemens Medical, Erlangen, Germany) (tube potential: 120 kVp, tube current: 58 mA, voxel size: 0.59 x 0.59 x 1.5 mm^3^) as well as a PET/MR measurement with the radioligand [^11^C]DASB on a Biograph mMR (Siemens Medical, Erlangen, Germany). A total calculated dose of 20 MBq/kg was delivered via a bolus and constant infusion protocol (1 min bolus, 179 min infusion). The 125-minute PET scan started 30-45 minutes after the initial bolus. During the PET examination, a T1-weighted magnetization-prepared rapid gradient-echo sequence (TE/TR = 4.21/3000 ms, voxel size 1 x 1 x 1.1 mm^3^) and the vendor-provided DIXON-VIBE sequence (TE1/TE2/TR = 1.23/2.46/3.60 ms, flip angle = 10, voxel size 2.6 x 2.6 x 3.1 mm^3^) were acquired. For the quantification of serotonin transporter binding, arterial blood samples were taken at minutes 120, 130, 140 post radioligand application.

### Attenuation correction approaches

A schematic overview for the creation of attenuation maps is shown in Figure 1. To implement the various pCT approaches, we curated an in-house database of MRI-CT pairs. For each subject in the database, the CT scan was co-registered to the corresponding T1-weighted MR image via a rigid transformation using SPM12. Images were resliced (4^th^ degree B-Spline) and proper alignment was visually verified. T1-weighted images were then normalized to MNI space (SPM12, default settings), using the tissue probability map (TPM) provided in SPM12. The head holder was masked using the TPM and background was set to -1024 HU. The resulting transformation was then applied to the co-registered CT. pCT generation followed the algorithms described in prior work, except for having CTs in normalised MNI space instead of individual space for the implementation of the modified MaxProb (mMaxProb) approach. For MaxProb (13,26), each CT was segmented into three tissue classes by intensity (I) thresholding: I < -500 HU (air), -500 HU ≤ I < 300 HU (soft tissue) and I ≥ 300 HU (bone) (13,30). For each voxel, the most prevalent tissue class at that location across all database CTs was determined. The pCT template was then generated by averaging Hounsfield units over voxels belonging to the most prevalent tissue class. The Boston method (12) differs from MaxProb, as implemented here, only in the averaging step: all voxels contribute to the voxel-wise mean, regardless of tissue class. For both methods, attenuation maps were scaled to linear attenuation coefficients (LAC) at 511 keV using bilinear scaling (31). For creation of Boston and mMaxProb pCTs, each subject’s T1-weighted image was normalized to MNI space, and the inverse transform was then applied to the template for transformation to individual subject space. Database CTs were first normalized to MNI space, ensuring that the only subject-specific deformation applied to the resulting attenuation maps was the inverse transformation back to native space. To improve computational efficiency, the MNI-space CTs were subsequently aggregated into template volumes. Accordingly, both the Boston method and the mMaxProb approach are classified as template-based in this manuscript.

**Figure 1:**
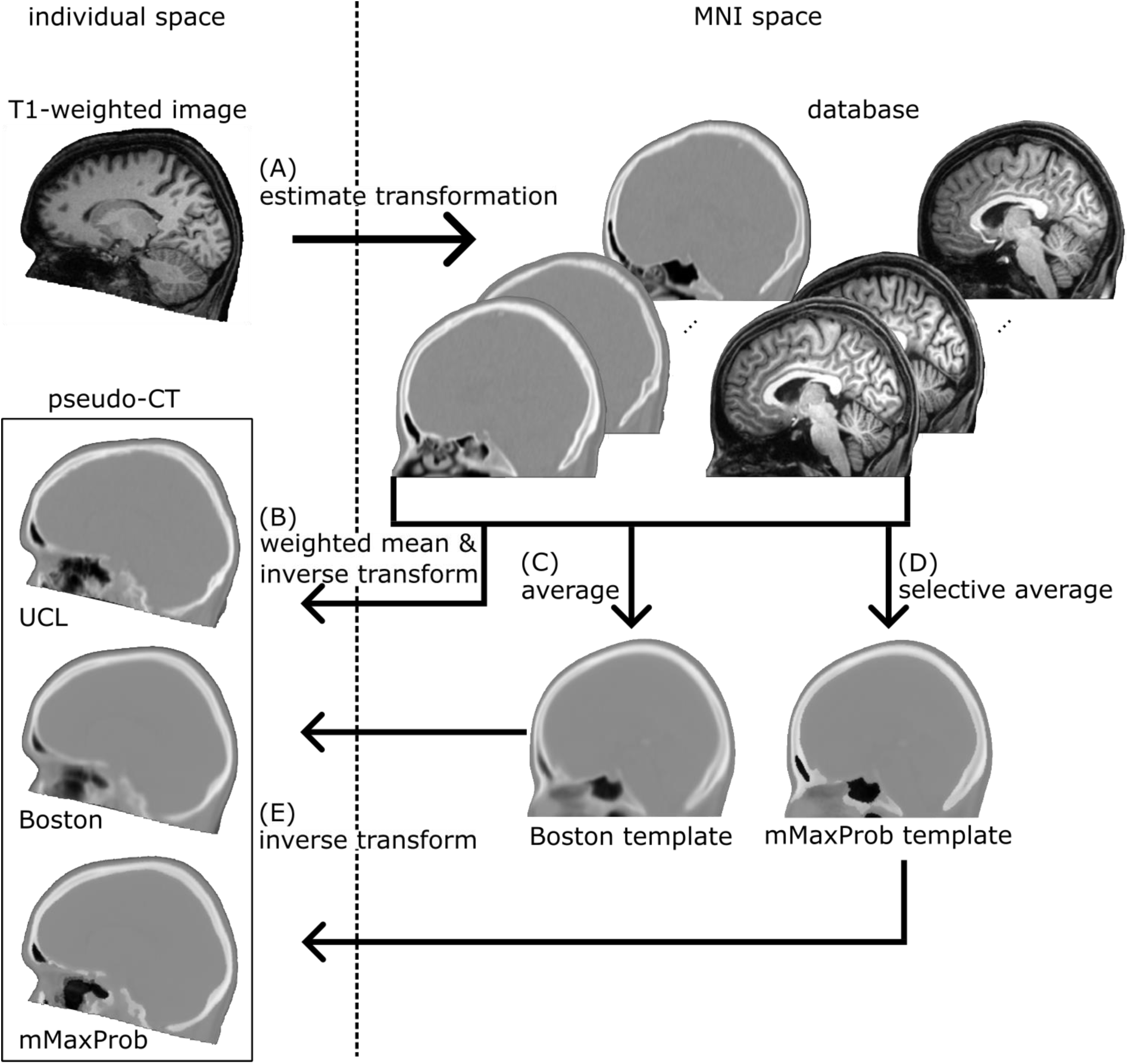
Schematic workflow for pseudo-CT (pCT) creation. The T1-weighted image of the subject for which a pCT should be created is provided in its native space (individual space). The database of CT and T1-weighted MR image pairs is provided in MNI space. The transformation of the T1-weighted image to MNI space is estimated first (A). UCL: a weighted mean of the database CTs based on local similarity between database and subject T1-weighted images is computed and transformed back to individual space (B). Boston: An average of database CT images yields a template (C). mMaxProb: Following segmentation, a tissue-selective average of database CT images yields a template (D). Templates are transformed back to individual space (E).

To investigate the effect of database size on performance, we created mMaxProb and Boston templates from subsets of 5, 10, 20, 40 and 60 subjects. Throughout the manuscript, the database size will be referred to as an index (e.g. mMaxProb_db20_). To minimize potential confounding effects, each subset was balanced for sex distribution and age range. In absence of an index referring to the specific database size, the modified implementation of MaxProb will be referred to as “mMaxProb” to distinguish it from the original implementation.

Additionally, the UCL approach (11) for pCT generation was adapted to work with the MNI space database. A convolution-based fast local normalized correlation coefficient (LNCC) (32) was calculated between a subject’s and each database T1-weighted image. Subsequently, LNCCs were used for ranking each voxel in the database images. Weights (W) were then calculated based on rankings (R) according to Eq1. Applying these weights to the database CTs yielded pCTs in MNI space. Pseudo-CTs were transformed to individual space in the same manner as for the template-based pCTs. This implementation of the UCL approach uses the full database size of 60 subjects. Unlike the MaxProb and Boston implementation, database CTs cannot be combined into a template, because the weighted mean of database CTs is subject-specific.

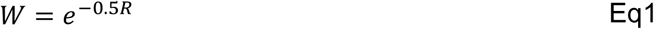

Since the field of view (FOV) of an image normalized in SPM12 does not cover the entire head and neck, we supplemented these pCTs with the vendor provided attenuation map (DIXON) (8), as described in (6). We intentionally did not replace the corresponding region in the CT, in order to capture a possible effect and to recreate a realistic use case. Finally, all pCTs were resliced (4^th^ degree B-Spline) to the DIXON attenuation map and converted to DICOM format for integration into the Siemens reconstruction pipeline. To preserve the metadata contained in the original DIXON attenuation map DICOM file, only image data was modified by using the dcmodify tool from the DICOM toolkit (DCMTK) package.

The three pCT approaches were validated in an independent sample, whose subjects were not part of the database. As a reference standard, attenuation maps were generated from each subject’s CT image (10,31). Analogous to the pCTs, CT scans were scaled to LACs at 511 keV, the headrest was masked, CTs were coregistered to subject’s T1-weighted image and resliced to match the dimensions of the DIXON attenuation map.

### Image Reconstruction

PET images were reconstructed using the Siemens reconstruction software (e7tools, Siemens Medical Solutions, Knoxville, USA) as described previously (33). Three consecutive 10-minute frames were reconstructed (ordinary Poisson-ordered subset expectation maximization, 3 iterations, 21 subsets, post-reconstruction filter 5 mm Hann, matrix size 344 x 344 x 127, voxel size 2.09 x 2.09 2.03 mm^3^, Zoom 1) in tracer equilibrium, starting 95 minutes after PET start. Attenuation correction was performed with each of the attenuation maps: CT, UCL, mMaxProb_db5-db60_ Boston _db5-db60_).

### Data processing and Quantification

PET data were processed using SPM12 with default parameters, unless otherwise specified. The three reconstructed PET frames were realigned (quality = 1, register to mean) to correct for head motion and then co-registered to the T1-weighted image. The normalization to MNI space was estimated with the T1-weighted image and then applied to the PET data. Imaging data was processed using SPM12 and MATLAB R2018b (The Mathworks Inc., Natick, MA, USA).

pCT performance was evaluated by quantifying serotonin transporter (SERT) binding from PET data, reconstructed with each attenuation map. Total volume of distribution (V_T_ = C_T_/C_P_) (1) was quantified using an equilibrium method (33–35). Accordingly, mean activity concentrations in radioligand equilibrium from the respective compartments were obtained for tissue (C_T_) and plasma (C_P_). Regional values were extracted for thalamus, hippocampus, amygdala, frontal lobe, temporal lobe, parietal lobe and occipital lobe, defined using the Harvard-Oxford atlas as provided with FSL (36), and an in-house atlas was used for cerebellar grey matter (37)

### Statistical Analysis

Performance of different attenuation correction approaches was analyzed in two steps. First, similarity between each pCT and the reference CT attenuation map was assessed using the Jaccard index (38). Separate indices were calculated for bone (LAC > 0.1cm^-1^) and soft tissue (0.05 cm^-1^ < LAC ≤ 0.1 cm^-1^) (7). Analyses were performed for both the full FOV as well as only the area covered by the original pCTs. Second, the effect of attenuation correction approaches on V_T_ was evaluated by mean relative error 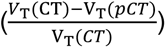 and mean absolute error relative to the CT-based attenuation correction. A repeated measures analysis of variance (rmANOVA) was performed for each ROI, to compare relative errors in V_T_ between the three pCT approaches. For Boston and mMaxProb, only the best performing database size was included. Bonferroni correction was used to correct for repeated testing (9 ROIs). Paired sample t-tests were used for post-hoc comparison in ROIs where the rmANOVA results were significant.

## RESULTS

A database of 60 MRI-CT pairs was curated and transformed to MNI space. Templates for mMaxProb and Boston pCTs were created for five database sizes (n = 60, 40, 20, 10, 5). The subcohorts did not differ significantly with regard to mean age (one-way ANOVA: p = 0.89, F = 0.275) or sex (chi-squared test: p = 0.98, χ²(4) = 0.39).

A visual comparison between CT and pCTs for a representative subject is shown in Figure 2. The template-based pCT Methods (Boston, mMaxProb) exhibited greater blurring at larger database sizes. In general, the greatest deviations from the reference CT were present in the posterior part of the skull and the paranasal sinuses. In these regions, mMaxProb pCTs tended to underestimate LAC values, ignoring the fine bone structures. Similarly, Boston and UCL failed to capture this complex combination of air, tissue and bone. UCL best captured variations within the skull.

**Figure 2:**
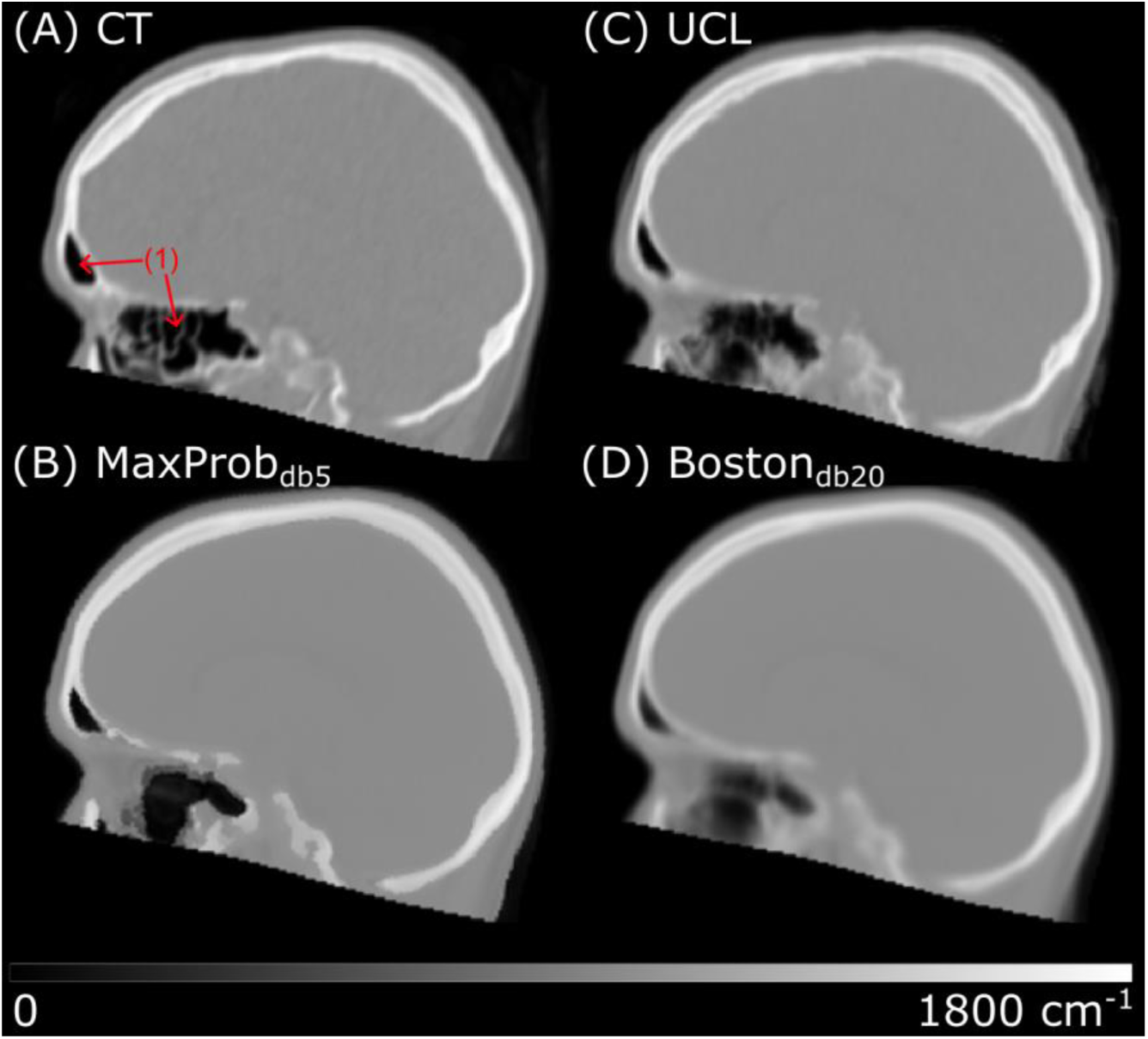
CT and pseudo-CT for a representative subject. The complex air-bone interface in the paranasal sinuses poses an issue in all pCT approaches. mMaxProb and UCL preserve sharp tissue borders whereas Boston exhibits obvious smoothing across tissues. Clear differences in size and tissue-bone composition are visible in the paranasal sinuses (1).

As assessed by the Jaccard index, similarity between pCTs in the segmentation of bone and soft tissue was comparable for the mMaxProb and Boston approach and consistent across database sizes, while the UCL pCTs performed numerically better, particularly for the bone segment (rmANOVA followed by pairwise comparison, all p < 0.001). As expected, the observed correspondence between pCT and CT was reduced when including the neck region that was supplemented by the DIXON attenuation map in the pCTs. A comprehensive overview of these results can be found in Table 2.

**Table 2:** Jaccard index (J) between computed tomography (CT) attenuation map and pseudo-CTs (pCTs). Similarity is evaluated for bone (LAC > 0.1cm^-1^) and soft tissue (0.05 cm^-1^ < LAC ≤ 0.1 cm^-1^) in the head region (H) without the neck and across the entire attenuation map (HN). Results are reported as mean ± SD.

| pCT | J(Bone<sub>H</sub>) | J(Bone<sub>HN</sub>) | J(Tissue<sub>H</sub>) | J(Tissue<sub>HN</sub>) |
| --- | --- | --- | --- | --- |
| UCL | 0.71 ± 0.07 | 0.58 ± 0.05 | 0.80 ± 0.03 | 0.72 ± 0.03 |
| MaxProb<sub>db60</sub> | 0.66 ± 0.07 | 0.54 ± 0.05 | 0.78 ± 0.03 | 0.72 ± 0.03 |
| MaxProb<sub>db40</sub> | 0.66 ± 0.07 | 0.54 ± 0.05 | 0.78 ± 0.03 | 0.72 ± 0.03 |
| MaxProb<sub>db20</sub> | 0.65 ± 0.06 | 0.53 ± 0.04 | 0.78 ± 0.03 | 0.72 ± 0.03 |
| MaxProb<sub>db10</sub> | 0.65 ± 0.06 | 0.53 ± 0.04 | 0.78 ± 0.03 | 0.72 ± 0.03 |
| MaxProb<sub>db5</sub> | 0.64 ± 0.06 | 0.53 ± 0.05 | 0.78 ± 0.03 | 0.72 ± 0.03 |
| Boston<sub>db60</sub> | 0.66 ± 0.07 | 0.55 ± 0.05 | 0.77 ± 0.03 | 0.71 ± 0.03 |
| Boston<sub>db40</sub> | 0.66 ± 0.07 | 0.55 ± 0.05 | 0.77 ± 0.03 | 0.71 ± 0.03 |
| Boston<sub>db20</sub> | 0.66 ± 0.06 | 0.54 ± 0.05 | 0.77 ± 0.03 | 0.71 ± 0.03 |
| Boston<sub>db10</sub> | 0.65 ± 0.06 | 0.54 ± 0.05 | 0.77 ± 0.03 | 0.71 ± 0.03 |
| Boston<sub>db5</sub> | 0.65 ± 0.06 | 0.53 ± 0.05 | 0.78 ± 0.03 | 0.72 ± 0.03 |

Mean relative error in SERT V_T_ (Table 3) was calculated between PET reconstructions using CT-based and pCT-based attenuation correction. Quantification errors were generally low and distributed around zero with a small negative bias in most ROIs, indicating an overestimation of V_T_ reconstructed with pCTs. Among the pCT approaches, UCL performed best, followed by mMaxProb_db5_ and Boston_db20_, all showing small relative mean error below 0.5% across the brain. No clear trend of database size and pCT performance was evident in the relative errors. Boxplots of mean relative quantification errors are provided in Figure 3 for the best performing pseudo-CTs (UCL, mMaxProb_db5_, Boston_db20_). Boxplots for all database sizes as well as mean absolute errors are provided in the supplement. The rmANOVA yielded significant results in amygdala, thalamus, cerebellar grey matter and parietal lobe. The post-hoc analysis showed significantly better performance of Boston_db20_ pCTs compared to mMaxProb_db5_ in amygdala (p < 0.001), thalamus (p < 0.001) and CGM (p < 0.001) and better performance in CGM than UCL (p = 0.02). mMaxProb_db5_ pCTs resulted in lower quantification errors in the parietal lobe compared to UCL (p = 0.04) and Boston_db20_ (p < 0.001). UCL performed better than mMaxProb_db5_ in CGM (p < 0.001). All tests were corrected for multiple testing using Bonferroni correction and were evaluated at a significance level of 0.05.

**Figure 3:**
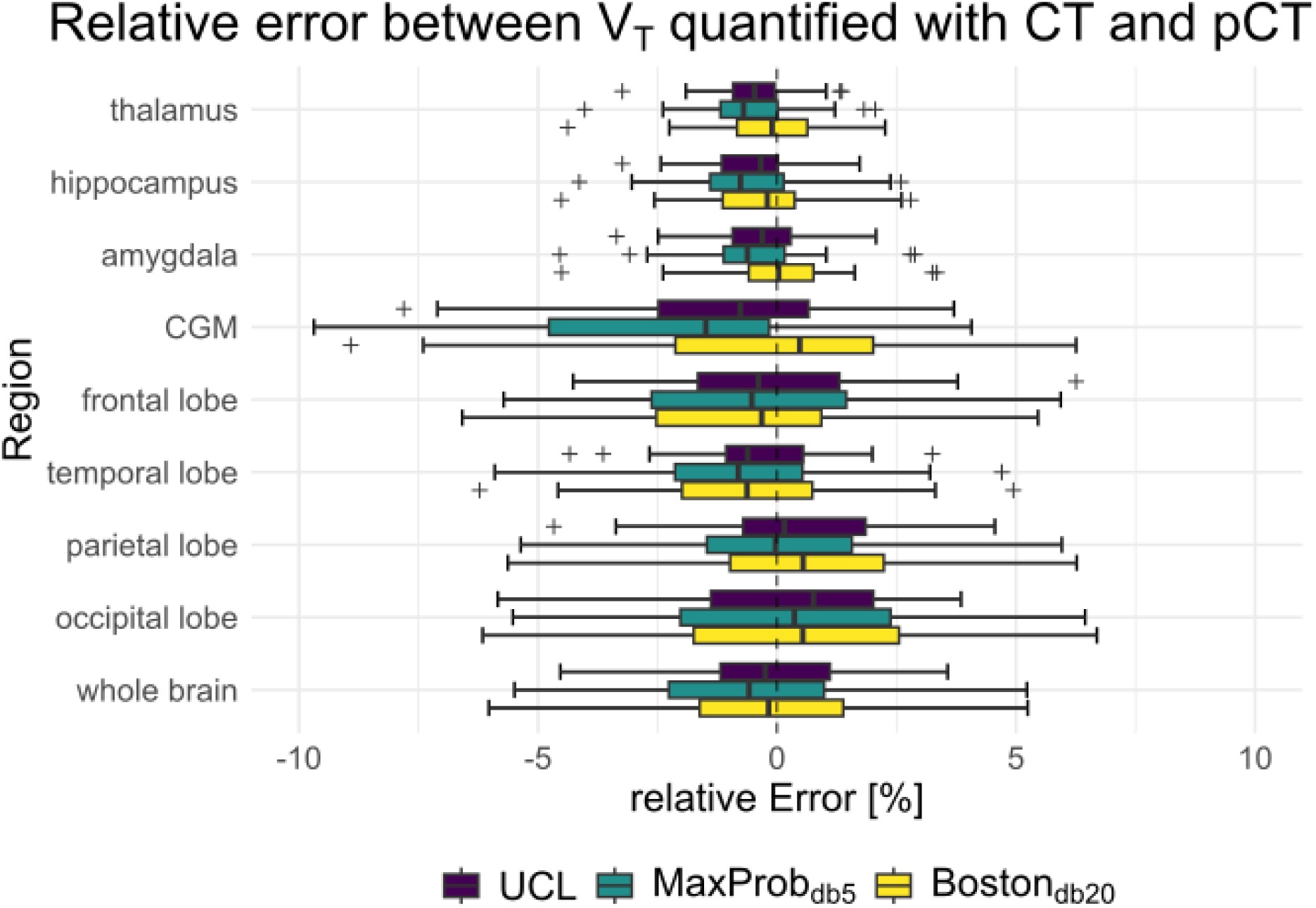
Boxplot showing mean relative quantification error 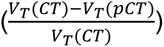 between PET reconstruction with CT and pCT. The three pCT methods UCL, Boston_db20_ and MaxProb_db5_ represent the best-performing pCT approaches. Cerebellar grey matter (CGM).

**Table 3:** Mean relative error 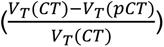 in V_T_ between PET reconstructed with CT or pseudo-CT for attenuation correction (mean ± SD in %). Bold font indicates the best-performing variants used in subsequent analyses.

|  | thalamus | hippocampus | amygdala | cerebellar<br>grey matter | frontal lobe | temporal<br>lobe | parietal lobe | occipital lobe | whole brain |
| --- | --- | --- | --- | --- | --- | --- | --- | --- | --- |
| **UCL** | **-0.48±1.05** | **-0.47±1.19** | **-0.29±1.25** | **-1.05±2.41** | **-0.06±1.70** | **-0.42±2.23** | **0.36±2.38** | **0.30±1.93** | **-0.03±2.77** |
| MaxProb<sub>db60</sub> | -1.10±1.36 | -1.21±1.49 | -0.95±1.55 | -1.95±2.87 | -1.34±2.29 | -1.60±2.68 | -0.75±2.92 | -0.93±2.37 | -1.19±3.65 |
| MaxProb<sub>db40</sub> | -1.28±1.32 | -1.36±1.47 | -1.06±1.52 | -2.23±2.90 | -1.46±2.27 | -1.79±2.70 | -0.87±2.92 | -1.20±2.38 | -1.35±3.56 |
| MaxProb<sub>db20</sub> | -0.74±1.33 | -0.95±1.43 | -0.58±1.54 | -0.94±2.84 | -0.99±2.26 | -1.30±2.69 | 0.18±2.94 | 0.14±2.35 | -0.60±3.64 |
| MaxProb<sub>db10</sub> | -1.05±1.36 | -1.07±1.47 | -0.90±1.55 | -1.64±2.91 | -1.11±2.26 | -1.31±2.70 | -0.92±2.94 | -0.51±2.40 | -1.00±3.58 |
| **MaxProb<sub>db5</sub>** | **-0.61±1.34** | **-0.66±1.49** | **-0.54±1.58** | **-2.27±2.92** | **-0.44±2.33** | **-0.91±2.71** | **-0.02±2.95** | **0.33±2.41** | **-0.35±3.62** |
| Boston<sub>db60</sub> | -1.03±1.50 | -1.13±1.60 | -0.86±1.65 | -1.78±2.97 | -1.50±2.38 | -1.46±2.81 | -0.83±2.99 | -1.00±2.48 | -1.24±3.71 |
| Boston<sub>db40</sub> | -1.10±1.44 | -1.15±1.57 | -0.88±1.59 | -1.92±2.90 | -1.52±2.36 | -1.52±2.71 | -0.86±2.96 | -1.18±2.41 | -1.30±3.74 |
| **Boston<sub>db20</sub>** | **-0.20±1.42** | **-0.34±1.52** | **-0.02±1.60** | **-0.25±2.93** | **-0.54±2.31** | **-0.58±2.79** | **0.54±3.07** | **0.58±2.46** | **-0.11±3.78** |
| Boston<sub>db10</sub> | -0.66±1.46 | -0.67±1.58 | -0.47±1.62 | -0.95±2.93 | -0.71±2.36 | -0.57±2.75 | -0.34±2.99 | -0.13±2.44 | -0.51±3.74 |
| Boston<sub>db5</sub> | -0.71±1.40 | -0.72±1.56 | -0.58±1.60 | -2.16±2.92 | -1.00±2.38 | -1.04±2.71 | -0.38±3.03 | 0.02±2.44 | -0.69±3.74 |

Processing time was reduced compared to the original algorithms by having the database and templates in MNI space. This reduced the number of spatial transformations per pCT to one, whereas in the original individual-space implementation of MaxProb and UCL, the number scales linearly with database size. When using the full database size, processing time is decreased by a factor of twenty. All three adapted pCT approaches perform similarly with regard to computational speed.

## DISCUSSION

This work adapts the MaxProb, Boston and UCL algorithms for pCT generation in MNI space and provides open access to the associated templates, database and code, to facilitate local implementation, ensure availability, and resolve legal barriers. The proposed approaches achieve comparable performance in the quantification of SERT V_T_ while substantially reducing computational demands compared to the original implementations.

In terms of segmentation overlap, UCL achieved the highest Jaccard index values for both bone and soft tissue, consistent with visual inspection of the attenuation maps. Since our SPM12-based pCTs do not cover the neck region, similarity to the reference CT was evaluated both for the entire head, supplemented with the DIXON attenuation map, and for the brain region alone. As expected, inclusion of the neck region reduced similarity scores, since the DIXON attenuation map lacks a bone component (8). The largest deviations in V_T_ were observed along the skull, where inaccuracies in LAC estimation can strongly impact quantification. The paranasal sinuses also posed a challenge, with prior reports noting variability in sinus size and composition (air vs. fluid) (26) and proposing region-specific masks to better capture the air–bone–soft tissue interface (10).

In general all three pCTs approaches yield low quantification errors (<0.5% whole brain average for UCL, mMaxProb_db5_, Bostondb_20_). Still, differences between pCTs were observed. These were small in magnitude but consistent across subjects as indicated by the rmANOVA. One of the sources for the observed differences stems from the averaging of the database in both template-based approaches. Specifically, Boston smoothed the entire image, whereas MaxProb only smoothed within tissue classes (bone, soft tissue or air), preserving sharp tissue boundaries. These differences arise from the underlying averaging strategies: Boston averages all database CT voxels, while MaxProb averages only within the most prevalent tissue class. Neither mMaxProb nor Boston capture inter-individual variability in bone contour variability as well as UCL. This is partly explained by the number of subjects contributing to each voxel: for MaxProb, at least one third of the database contributes, whereas for Boston, all subjects contribute equally. UCL pCTs are less blurred due to the exponential weighting, and larger databases should further improve performance as more individual variation is captured. In contrast larger databases increase blurring in Boston and MaxProb pCTs. It is noteworthy that the re-implementation of the MaxProb method within MNI space introduces methodological deviations from the original formulation. The initial approach entails direct registration of database MRI-CT pairs to native subject space, with subsequent label fusion performed in that same space, thus yielding subject-specific pCTs. The use of multiple independent registrations in the multi-atlas-based original approach enhances robustness (39).

In comparison to the original reports (11,12,26), our results are broadly consistent with previously observed performance trends for MaxProb, Boston, and UCL methods. Our results show UCL achieving the highest Jaccard indices and lowest V_T_ errors. Moreover, (13) also demonstrated that the improvement in performance by increasing database size quickly levels out, which is consistent with our results. Quantitative accuracy in our study, low bias and small relative error compared with CT-based AC, are also comparable to that reported in a large multi-centre evaluation (6), supporting the generalizability of these methods when adapted to MNI space. Notably, a spatially normalized database has several advantages and aligns with the FAIR principles. First, it achieved comparable performance at substantially reduced computational cost. Second, the database resolves data protection limitations inherent to online-only tools such as the original UCL implementation operating with T1-weighted images in native space where original faces may be reconstructed (40,41). That is, the non-linear spatial transformations applied during database generation make re-identification of subjects unlikely as transformation parameters are not publicly available. Third, this enables making the entire database openly accessible, which in turn promotes open software solutions and guarantees availability as the database may be stored locally. Fourth, independence from the provider enhances reproducibility of data analyses.

### Limitations

Our implementation was validated using [^11^C]DASB dynamic PET data. While we acknowledge the diversity of PET radioligands, it is important to note that these methods are adaptations of established methods and the original publications show similar results and have since been replicated (6), emphasizing robustness across diverse datasets. While structural brain alterations have been reported in MDD (42,43), expected variations in LACs at 511 keV (44) are not anticipated to significantly impact AC. The spatial coverage of the pseudo-CT templates corresponds to that of the SPM12 tissue probability map, ending below the nose and excluding the neck region. This region was not replaced with the DIXON attenuation map in the CT reference standard, yet performance remained acceptable. We also note that scanner-integrated DIXON-based pCT generation tools (45), which were unavailable for our dataset, could further enhance the presented methods by supplementing the template-based attenuation map with more accurate subject-specific attenuation information. Dense hair (braids, dreadlocks) is not accounted for in all pCT approaches and has been shown to introduce localized errors in the PET reconstruction (46). As with all database AC methods, accurate spatial normalization is a prerequisite. These approaches may perform poorly in subjects with major structural abnormalities (e.g., lesions, malformations).

## Conclusion

We present an openly available database of CT-T1 image pairs in MNI space, associated templates, and a streamlined pCT generation pipeline for PET/MRI attenuation correction. This approach resolves key data protection concerns, minimizes computational costs, and facilitates integration into custom reconstruction workflows. The local installation mode ensures long-term accessibility and adaptability to evolving research needs.

Beyond PET/MRI, the pseudo-CT tool introduced in this work may support other neuroimaging applications, such as planning functional transcranial ultrasound stimulation (47), as well as simulations for transcranial magnetic stimulation (48,49). Future developments should incorporate subject-specific information, such as implants or EEG electrode positions, to further enhance accuracy.

## ACKNOWLEDGEMENTS

We thank the graduated team members and the diploma students of the Neuroimaging Lab (NIL, headed by R. Lanzenberger) as well as the clinical colleagues from the Department of Psychiatry and Psychotherapy of the Medical University of Vienna for clinical and/or administrative support. In details, we would like to thank S. Kasper, D. Rujescu for their medical support, L. Rischka, G.M. James for technical support, N. Berroterán-Infante, W. Wadsak, M. Mitterhauser for administrative support and C. Vraka and C. Philippe for radioligand synthesis.

## AUTHOR CONTRIBUTIONS

Study design R.L., A.H.

Data acquisition M.M., L.R.S., G.M.G., G.G., M.B.R.

Methods C.M., A.H., M.B.R.

Data Analysis C.M.

Manuscript preparation – Writing C.M., A.H., M.B.R

Manuscript preparation - Reviewing C.M., M.M., I.M., L.R.S., L.N., G.M.G., G.G.,

M.H., N.C., A.HAM., R.L., A.H., M.B.R.

## STATEMENTS AND DECLARATIONS

### Ethical considerations

The study was approved by the Ethics Committee of the Medical University of Vienna (1307/2014) and procedures were carried out in accordance with the Declaration of Helsinki.

### Consent to participate

All subjects gave written informed consent after detailed explanation of the study procedures.

### Consent for publication

Not applicable.

### Data availability

Raw data will not be publicly available due to reasons of data protection. Processed data and custom code can be obtained from the corresponding author with a data-sharing agreement, approved by the departments of legal affairs and data clearing of the Medical University of Vienna. The Matlab code for creation of pseudo-CTs is available at https://github.com/NeuroimagingLabsMUV/Milz_2025_pseudoCT under a CC BY-NC-SA license. Imaging data, including templates and individual CTs and T1-weighted MRI scans in MNI space, are available at https://doi.org/10.5281/zenodo.17953384 under a CC BY-NC-NDlicense.

