## Supplemental Figures and Tables for "Open-access template and database approaches for pseudo-CT generation in brain PET/MRI attenuation correction"

*Running title:*

*Template-Based Attenuation Correction in PET/MRI*

Christian Milz[1,2], Matej Murgaš[1,2], Inés Merida[3,4], Leo R Silberbauer[1,2], Lukas Nics[5], Godber M Godbersen[1,2], Gregor Gryglewski[1,2], Marcus Hacker[5], Nicolas Costes[3,4], Alexander Hammers[6,7], Rupert Lanzenberger[1,2], Andreas Hahn[1,2,#], Murray B Reed[1,2]

*[1]Department of Psychiatry and Psychotherapy, Medical University of Vienna, Austria*
*[2]Comprehensive Center for Clinical Neurosciences and Mental Health, Medical University of Vienna, Austria*
*[3]Centre de Recherche en Neurosciences de Lyon, Université Claude Bernard Lyon 1, France*
*[4]CERMEP-Imagerie Du Vivant, Lyon, France*
*[5]Department of Biomedical Imaging and Image-guided Therapy, Division of Nuclear Medicine, Medical University of Vienna, Austria*
*[6]School of Biomedical Engineering and Imaging Sciences, King's College London, UK*
*[7]King's College London & Guy's and St Thomas' PET Centre, St Thomas' Hospital, London, UK*

| | thalamus | hippocampus | amygdala | cerebellar grey matter | frontal lobe | temporal lobe | parietal lobe | occipital lobe | whole brain |
|---|---|---|---|---|---|---|---|---|---|
| UCL | 1.26±0.99 | 1.42±1.20 | 1.27±1.13 | 2.85±1.40 | 2.30±1.14 | 2.61±1.36 | 3.10±1.13 | 2.58±1.10 | 2.87±2.01 |
| $MaxProb_{db60}$ | 1.73±1.18 | 1.91±1.36 | 1.71±1.29 | 3.67±1.52 | 3.10±1.49 | 3.04±1.53 | 3.69±1.38 | 3.24±1.25 | 3.75±2.52 |
| $MaxProb_{db40}$ | 1.82±1.18 | 1.99±1.36 | 1.75±1.27 | 3.77±1.53 | 3.17±1.51 | 3.09±1.54 | 3.75±1.43 | 3.32±1.25 | 3.81±2.51 |
| $MaxProb_{db20}$ | 1.45±0.86 | 1.59±0.92 | 1.44±0.95 | 3.36±1.34 | 2.84±1.24 | 2.77±1.24 | 3.54±1.05 | 2.98±0.94 | 3.36±2.06 |
| $MaxProb_{db10}$ | 1.70±1.20 | 1.83±1.36 | 1.68±1.31 | 3.62±1.48 | 3.02±1.44 | 3.08±1.60 | 3.71±1.31 | 3.21±1.25 | 3.66±2.41 |
| $MaxProb_{db5}$ | 1.56±1.08 | 1.73±1.30 | 1.59±1.26 | 3.57±1.43 | 3.04±1.40 | 3.04±1.51 | 3.75±1.23 | 3.19±1.18 | 3.90±2.55 |
| $Boston_{db60}$ | 1.74±1.26 | 1.92±1.41 | 1.73±1.34 | 3.77±1.58 | 3.09±1.51 | 3.09±1.64 | 3.69±1.48 | 3.29±1.33 | 3.75±2.49 |
| $Boston_{db40}$ | 1.75±1.25 | 1.91±1.41 | 1.69±1.32 | 3.77±1.57 | 3.11±1.52 | 3.05±1.58 | 3.70±1.53 | 3.29±1.32 | 3.76±2.60 |
| $Boston_{db20}$ | 1.36±0.83 | 1.45±0.87 | 1.40±0.96 | 3.32±1.35 | 2.71±1.16 | 2.83±1.35 | 3.60±1.14 | 2.95±0.99 | 3.40±2.02 |
| $Boston_{db10}$ | 1.58±1.18 | 1.75±1.33 | 1.59±1.28 | 3.54±1.46 | 2.93±1.35 | 3.00±1.59 | 3.67±1.29 | 3.14±1.23 | 3.61±2.36 |
| $Boston_{db5}$ | 1.58±1.18 | 1.76±1.36 | 1.60±1.31 | 3.66±1.47 | 3.02±1.46 | 3.03±1.53 | 3.75±1.25 | 3.22±1.22 | 3.88±2.63 |

Supplementary Table 1: Mean absolute relative error ($|\frac{V_T(CT)-V_T(pCT)}{V_T(CT)}|$) in $V_T$ between PET reconstructed with CT or pseudo-CT for attenuation correction (mean ± SD in %).

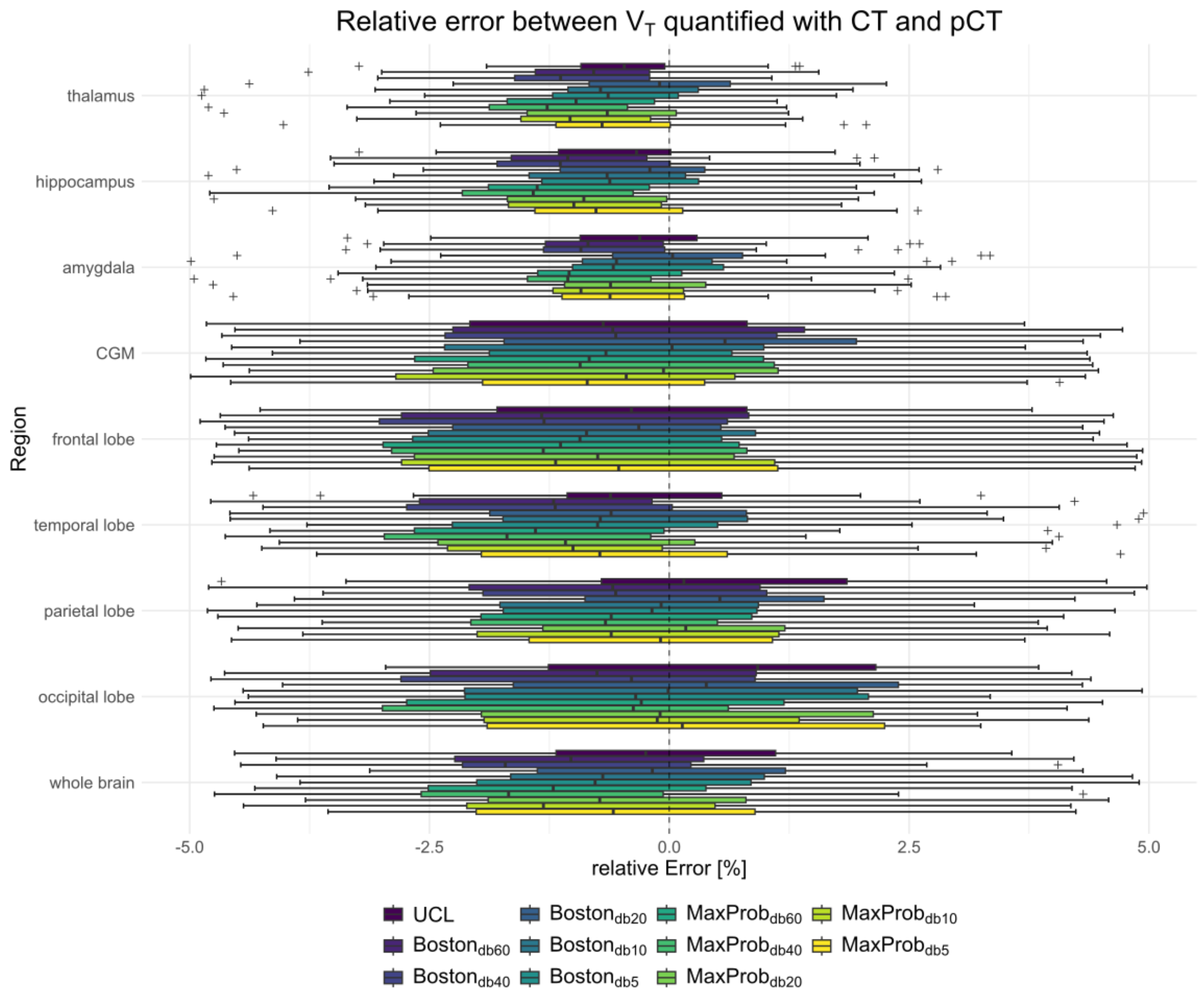


Supplementary Figure 1: Boxplot showing mean relative quantification error ($\frac{V_T(CT)-V_T(pCT)}{V_T(CT)}$) between PET reconstruction with CT and pseudo CT methods. Cerebellar grey matter (CGM).

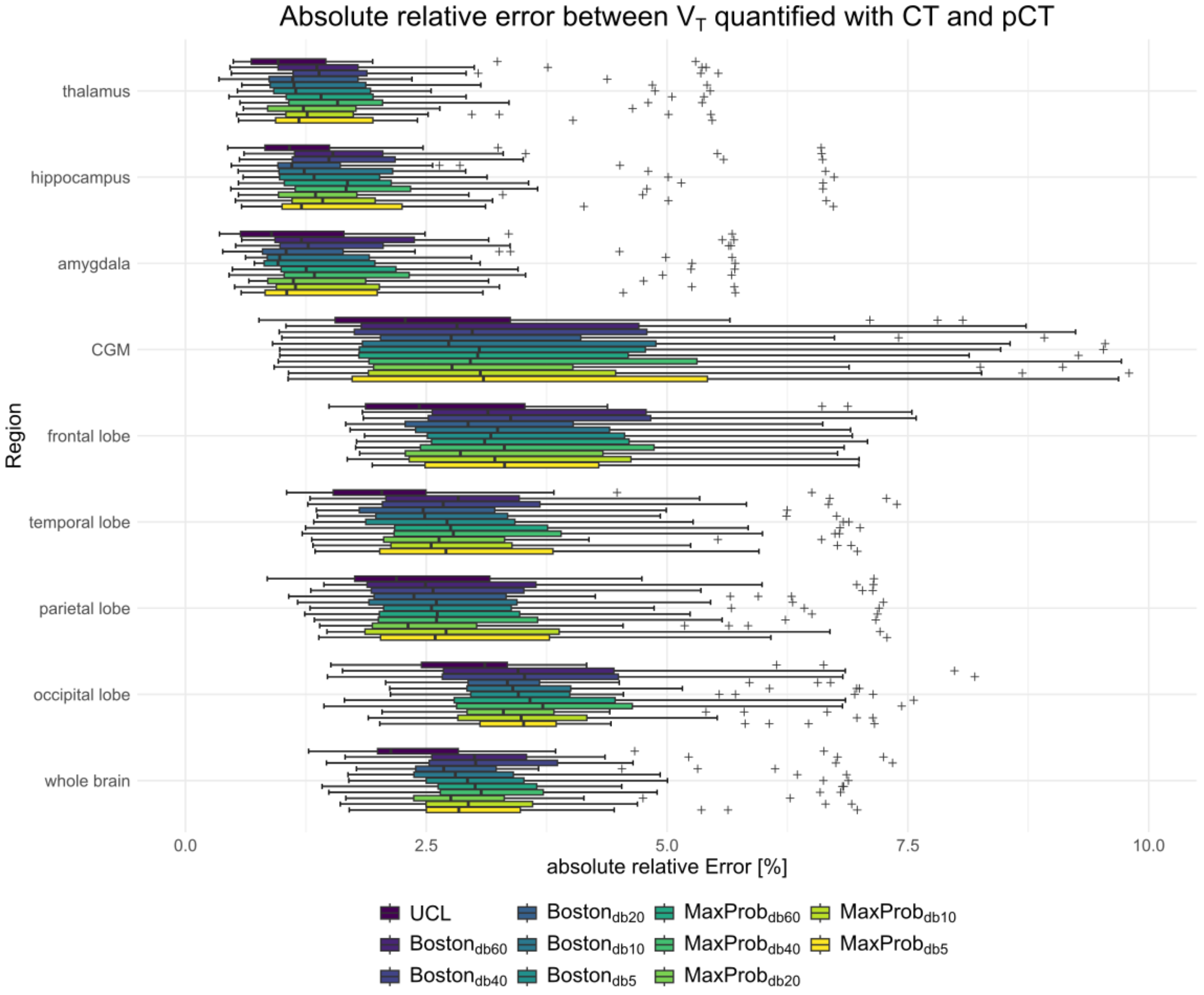


Supplementary Figure 2: Boxplot showing mean absolute relative quantification ($\left|\frac{V_T(CT)-V_T(pCT)}{V_T(CT)}\right|$) error between PET reconstruction with CT and pseudo CT methods. Cerebellar grey matter (CGM)
